# Sexual imprinting overrides order effects

**DOI:** 10.1101/828236

**Authors:** Luis Moreira, Léa Zinck, Kensaku Nomoto, Susana Q. Lima

## Abstract

Mate choice is a complex decision that requires the integration of cues from potential mates with individual preferences. Choosers’ preferences are shaped by recent events, early life experience and by the evolutionary history of its own species. To better understand the interaction between these factors, we studied mate choice in the female house mouse, *Mus musculus*. Females of one of the *musculus* subspecies, *Mus musculus musculus*, show preference for males of their own subspecies compared to males of the sibling subspecies, *Mus musculus domesticus*. Such an assortative preference is ecologically relevant at contact zones, where it contributes to the reproductive isolation of sympatric populations and can be reproduced in controlled laboratory conditions, but its origins are still under debate. Here, we show that female mouse mate choice depends on both early postnatal life experience and the order of prospective mates encountered as an adult and that these effects interact asymmetrically. Whereas females raised in their normal *M. m. musculus* environment display a robust assortative preference, females fostered in a *M. m. domesticus* family prefer the first male encountered, regardless of subspecies. Thus, early life experience of *M. m. musculus* females, when concordant with genetic self-identity, overrides sampling order effects, ensuring robust assortative choice. In the absence of this match between phylogeny and early life experience, first impression effects dominate mate choice.

---

Assortative mate choice is one of the mechanisms underlying the reproductive isolation between the two subspecies of house mouse, *M. m. musculus* and *M. m. domesticus.* These two lineages diverged from a common Asian ancestor which followed different human migratory routes and have subsequently established a secondary contact zone in central Europe [1], where they exhibit asymmetric mate preference: *M. m. musculus* females prefer their own subspecies, whereas *M. m. domesticus* females mate indiscriminately with both [2]. Using inbred strains of wild-derived mice as representatives of the *M. m. musculus* subspecies and laboratory mice as representatives of *M. m. domesticus*, the preference of *M. m. musculus* females can be studied under laboratory conditions, where the social preference exhibited in a limited-contact paradigm is highly correlated with mating decisions [3].

To better understand the mechanisms underlying the mate choice of *M. m. musculus* females, we investigated the role played by early life experience in the establishment of the assortative preference. To do so, we fostered *M. m. musculus* females in a *M. m. domesticus* family at birth and assessed their mate preference during adulthood (Figure 1A and B, *M. m. musculus* ♂◻ = MUS, *M. m. domesticus* ♂◻ = DOM). We found that adoption disrupts the typical assortative preference exhibited by control females (Figure 1C and D, Test1 MUS preference), suggesting that mate preference in *M. m. musculus* females is strongly influenced by the early life experience of the female. To assess the stability of mate choice, we retested females two days later (Test2). Control females showed increased overall assortative preference (Figure 1C, Test2 MUS preference), while adopted females maintained no preference at the population level (Figure 1D, Test2 MUS preference). However, closer inspection of the behaviour of adopted females, revealed instead a widespread distribution in terms of male preference (Figure 1C). While some females spent more time with the MUS males, others preferred DOM males, and females increased their preference for the initially chosen male from Test1 to Test2. *M. m. musculus* females fostered in a non-familiar *M. m. musculus* environment at birth exhibited the typical assortative preference (Supp Figure 1A), indicating that pup handling and/or nest disturbance performed during the adoption manipulation do not disrupt female preference. Fostered and control females also showed similar behaviour during the habituation periods, including similar latencies of first entry into the compartment of the male (Supp Figure 1B, C, D). It is therefore unlikely that the disruption in preference was caused by anxiety due to differences in maternal behaviour [4].

**Figure 1.**
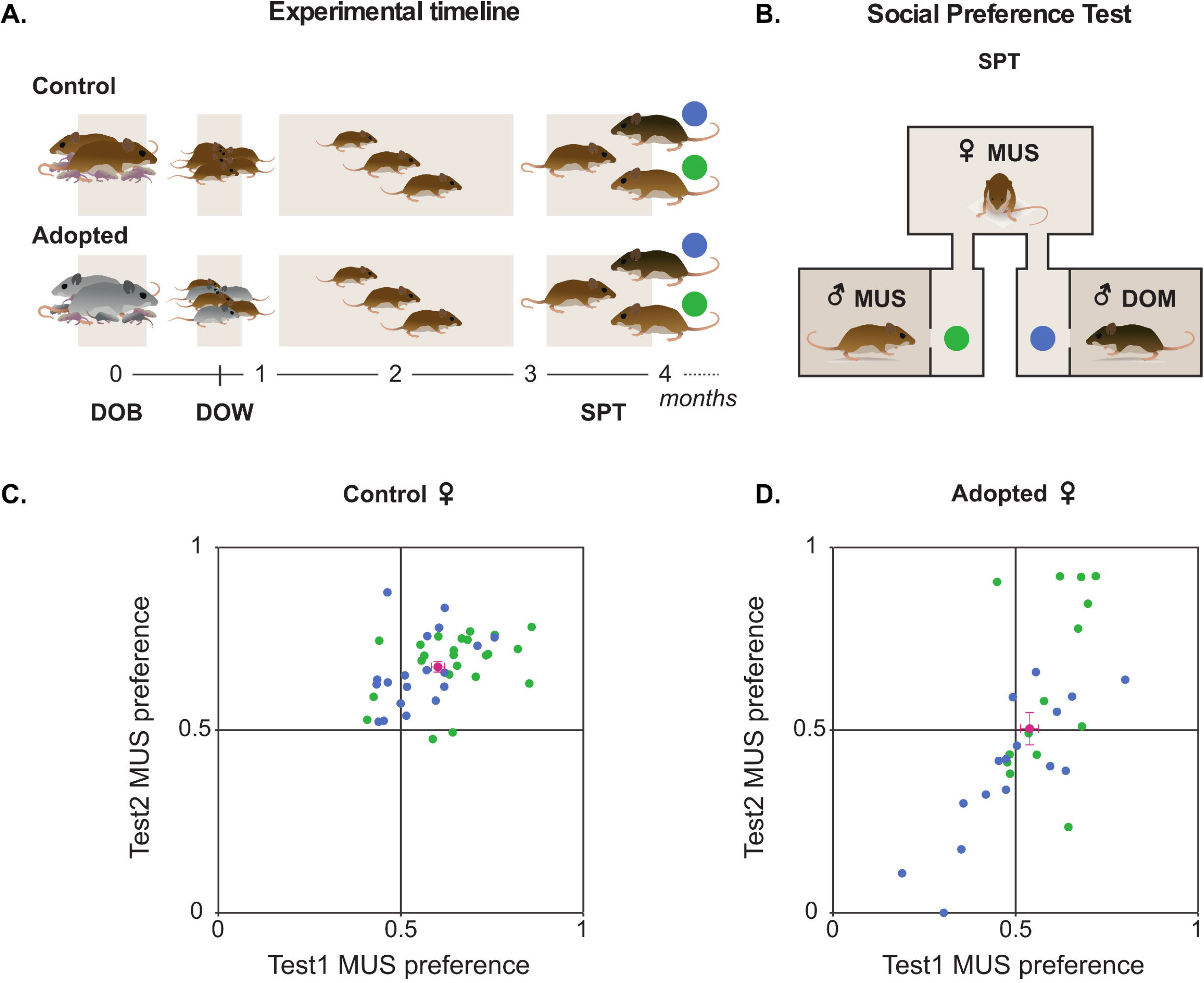
Mate choice of *M. m. musculus* females depends on early life experience and the order of exposure to males during adulthood. **(A)** *M. m. musculus* females were raised in a *M m. musculus* (control, n = 42) or a *M. m. domesticus* (adopted, n = 30) family from birth (DOB) until weaning (DOW). **(B)** Adult females’ mate preference was assessed in a social preference test (SPT) at around 3-4 months of age. T he initial preference was assessed during the first SPT trial (Testl) and the stability of the preference was examined during the second SPT (Test2). As representatives of the *M. m. musculus* subspecies we used the wild-derived strains of mice PWK and PWD, and as representatives of the *M. m. domesticus* we used C57BL/6 and B alb/c (please see methods for details and [3]). **(C)** Preference scores of control females were calculated as the time spent with the *musculus* male divided by the time spent with both males (green and blue dots are individual data, and in magenta is the mean ± standard error of the mean (SEM); green dots represent females that visited MUS males first, and blue dots represent females that visited DOM males first). **(D)** Same as (C) but for adopted ales.

Although adoption of *M. m. musculus* females in a *M. m. domesticus* environment disrupts female preference, indicating that the natural assortative choice does not rely exclusively on genetic/self-referencing matching mechanisms [5], it does not switch it to a preference for DOM either (as it has been shown using laboratory strains of mice [6]). This result suggests that, despite a role for sexual imprinting in normal conditions, imprinting on the foster *M. m. domesticus* family did not take place or that the adopted females may have learned to be non-choosy as *M. m. domesticus* females [2], and another local decision process might play a role in the choice of adopted females. As the behaviour was a “free-choice” test in a Y-maze and the first encounter was determined by whatever sequence the female happened to enter the arms in, we performed logistic regression analysis to test if the first male encountered could predict female preference (despite the fact that the first entry was random for both groups of females, *X*^2^ test, control: p = 0.82; adopted: p = 0.90).

Here we show two models of preference score for MUS males, the first for Test1 and the second for Test2. In the first model we have preference for MUS male as a function of early life experience (Control or Adopted), the identity of the first male visited during Test1 (MUS or DOM), the duration of this visit (standardized), as well as the interaction between these last two variables. For Test2 we have preference for MUS as a function of the early life experience, preference during Test1, as well as the interaction between the two terms. The model for Test1 shows that:
Predicted logit of (Test1_MUS_Pref) = 0.60 + (−0.27)*Adopted + (−0.64)*DOM + (0.11)*1ST_Visit_Duration + (1)*DOM*1ST_Visit_Duration.

According to the logistic regression model, these predictors explain 28% of the variance (R^2^ = 0.28), with preference for MUS in Test1 negatively related to being adopted (p < 0.05) and visiting DOM first (p < 0.001), with no significant effect of the duration of the first visit (p = 0.76), although a trend exists between the visit duration and the first male visited when this one was DOM (p = 0.06). In addition, the odds of an adopted female preferring DOM are 1.31 (1/e^−0.27^) times greater than the odds of a control female, and the odds of any female that visited DOM first preferring DOM are 1.90 (1/e^−0.64^) times greater than a female that visited MUS first. Therefore, adopted females are less likely to prefer MUS compared to controls, and visiting DOM first decreases the probability of either type of females preferring MUS.

On the other hand, the model for Test2 showed that:
Predicted logit of (Test2_MUS_Pref) = −0.01 + (−5.16)*Adopted + (1.09)*Test1_MUS_Pref + (8.35)*Adopted*Test1_MUS_Pref.

These predictors explained 33% of the variance (R^2^ = 0.33) showing that the preference for MUS in Test2 is negatively related to being adopted (p < 0.001), with no significant effect of preference during Test1 (p = 0.25), except for the interaction term, showing that in adopted females Test2 preference was positively related to the preference scores during Test1 (p < 0.001). The odds of an adopted female preferring DOM on Test2 were 173.39 (1/e^−5.16^) times greater than a control female, most likely dependent on familiarization processes that occurred during Test1. Thus, as in Test1, females adopted into a *M. m. domesticus* family are less likely to prefer MUS males, and the higher the preference in Test1 the more likely it is that an adopted female prefers the same male during Test2.

Studying mate choice in a biologically relevant context suggests that female choice in the house mouse depends on the interaction between at least two kinds of experience: exposure to concordant subspecies-specific cues during early life and the identity of the first male encountered during adulthood. Our adoption experiments revealed that if normal sexual imprinting is disrupted, the order of exposure to putative mates dictates female choice. Serial position and first impressions effects, which are well-documented cognitive biases in humans [7], but mostly unexplored in other species (but see [8] for influence of first visit on choice in mice), are likely to affect female choice since exposure to potential mates is done sequentially. Exposure order is particularly influential in contact zones where there is a high probability of females encountering males from a sibling species first. Sexual imprinting, which in the case of the house mouse appears to depend on the concordance between female and family species-specific cues [9], may override positional effects during female sampling. Since mice exhibit parental care and youngsters are normally raised in a family of their own subspecies, imprinting on family cues can provide a strong reproductive barrier that ensures assortative mate choice. This asymmetric interaction between mechanisms is likely to contribute to the strong reproductive isolation occurring at the *M. m. musculus* / *domesticus* contact zone.

## Materials and Methods

### Animals

C57BL/6J, BALB/cJ, PWD/PhJ and PWK/PhJ mice were ordered from The Jackson Laboratories and maintained in our animal facility. Animals were weaned at 21 days and housed in same-sex groups in stand-alone cages (1284L, Techniplast, 365 × 207 × 140 mm) with access to food and water ad libitum. To enhance the female estrous cycle [10], male-soiled bedding or urine was placed regularly in female home cage as described previously [3]. Mice were maintained on a 12:12 light/dark cycle and experiments were performed during the dark phase of the cycle, under red dim light. In all the experiments, we studied the preference of *Mus musculus musculus* PWD/PhJ females (with different experiences from birth to weaning) for PWK/PhJ males of the *M. m. musculus* subspecies and *M. m. domesticus* males (either C57BL/6J) using social-preference tests (SPT). All experiments were approved by the Animal Care and Users Committee of the Champalimaud Neuroscience Program and the Portuguese National Authority for Animal Health (Direcção Geral de Veterinária).

### Adoption experiments

Breeding pairs of mice of PWD/PhJ and BALB/cJ strains were started at the same time to be synchronized in order for us to adopt PWD/PhJ littermate females on the day of their birth, in a recipient breeding containing recently born pups (two-days old maximum). The recipient breeding was either of the BALB/cJ strain to study the role of female early experience on mate preference (adopted group), or of the PWD/PhJ strain to control for adoption manipulation (control adoption group). Moreover, in order to control for the overall number of pups in the recipient breeding, the female (and not the male) pups of the recipient breedings were removed and replaced by the adopted PWD/PhJ female pups (without their brothers). Every adopted PWD/PhJ female was thus in contact with her adopted mother, father and brothers, as well as with her blood PWD/PhJ littermate sisters. At 21 days, littermate adopted PWD/PhJ females were weaned and housed together until testing. No adoption has failed following this procedure.

### Social-preference tests

When the females of the different treatment groups reached 4 months of age, social-preference tests (SPT) were conducted as described in [3] to investigate to what extent mate choice is influenced by early experience. Briefly, the behavioural apparatus consisted of three transparent acrylic boxes (200 × 150 × 150 mm) connected by acrylic tubing to allow the subject female to go freely from the center box to the left and right boxes containing the stimulus males. Plastic partitions divided the male’s box to limit contact between the male and female but allowing facial contact and exchange of both volatile and non-volatile scents between the animals. The females were habituated for 15 minutes to the behavioural apparatus, without males, on three consecutive days (Hab1-3 on Day1-3). Then, on Day4 and Day6 after a similar habituation period, preference of the females for a *musculus* over a *domesticus* male was assessed during a 15-minute test (Hab4+Test1 on Day4; and Hab5+Test2 on Day6). Males were randomly assigned to the left or right box, in a balanced manner, and this assignment was kept constant for the same female between Test1 and Test2. After the tests were finished, we performed vaginal smears to determine the estrous state of the females, and we confirmed that the estrous state did not affect female preference in the SPT as described in [3] (data not shown). Sexually naïve *musculus* PWD/PhJ females (adopted or not) were always used as test subject, and sexually experienced males as stimulus. PWK/PhJ males were used as representative of the *musculus* subspecies while C57BL/6J males were used as representative of the *domesticus* subspecies.

### Behavioural analysis

Video recording was performed using a Sony camera (HDR HC7E) connected to a computer running Virtual Dub software to acquire images. Off-line analysis of videos was performed using Bonsai [11] and the behaviour of the female was analysed using MATLAB. The latency of first entry, the number of entries and re-entries (when the female returns to the same male’s compartment) into the male’s compartment, as well as the time spent in each male box were scored. Females’ preference score for *musculus* males was calculated as (amount of time spent with *musculus* male) / (total amount of time spent with the two males).

### Statistical analysis

Chi-squared tests were performed using MATLAB. To test the effects of adoption and first visit on the preference of MUS females we performed logistic regression analysis using Rstudio software (“betareg” package). For Test1, the model included the following terms: a) dependent variable: preference for *musculus* male; b) fixed factors: adoption treatment (control = 0; adopted = 1), identity of the first male visited (DOM = 0; MUS = 1) and duration of the first visit (standardized; continuous variable); c) interaction term between the identity of the first male visited and the duration of the first visit. Other interaction terms were removed from the analysis when not significant. For Test2, the model included the following terms: a) dependent variable: preference for *musculus* male; b) fixed factors: adoption treatment and preference during Test1 (continuous); c) interaction term between the adoption treatment and preference during Test1.

---

**Figure S1.**
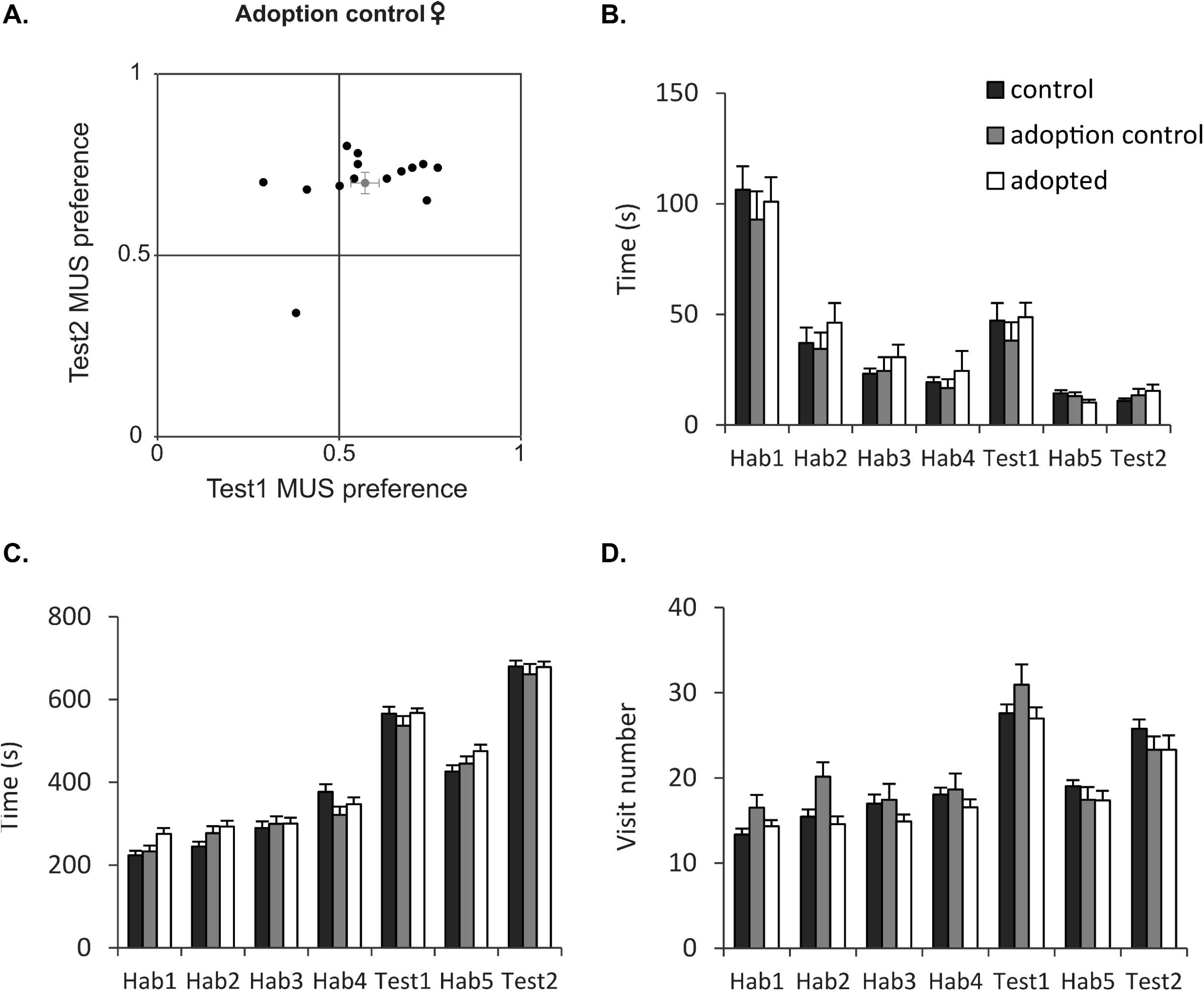
Adopted and control females exhibit similar behaviour. **(A)** Females adopted in a non-familiar *musculus* breeding at birth showed a normal assortative preference in both Testl and Test2. Control (unmanipulated; black), adoption control (fostered in *M. m. musculus* family; grey) and adopted females (fostered in *M. m. domesticus* family; white) showed **(B)** similar latency of first entry, **(C)** similar time spent with both males, and **(D)** similar number of visits to the male boxes during the habituation (Habl, Hab2, Hab3, Hab4, Hab5) and test (Testl, Test2) periods.

